# *Vibrio cholerae* Arrests Intestinal Epithelial Proliferation through T6SS-dependent Activation of the Bone Morphogenetic Protein Pathway

**DOI:** 10.1101/2023.06.29.547108

**Authors:** Xinyue Xu, Edan Foley

## Abstract

To maintain an effective barrier, intestinal epithelial progenitor cells must divide at a rate that matches the loss of dead and dying cells. Otherwise, epithelial breaches expose the host to systemic infection by gut-resident microbes. Unlike most pathogens, *Vibrio cholerae* blocks tissue repair by arresting progenitor proliferation in the *Drosophila* infection model. At present, we do not understand how *Vibrio* circumvents such a critical antibacterial defense. In a series of genetic experiments, we found that *V. cholerae* blocks epithelial repair by activating the growth inhibitor Bone Morphogenetic Protein (BMP) pathway in progenitors. Specifically, we discovered that interactions between *Vibrio* and gut commensals initiate BMP signaling via host innate immune defenses. Notably, we found that *Vibrio* also activates BMP and arrests proliferation in zebrafish intestines, indicating an evolutionarily conserved link between infection, BMP and failure in tissue repair. Combined, our study highlights how enteric pathogens engage host immune and growth regulatory pathways to disrupt intestinal epithelial repair.

## INTRODUCTION

To maintain an effective barrier against harmful environmental agents, multipotent Intestinal Stem Cells (ISCs) divide at a pace that matches the loss of dying epithelial cells. For example, infectious microbes frequently induce host defenses that prompt the expulsion of damaged epithelial cells ^1–5^. In response, ISCs divide at an accelerated rate to ensure an adequate supply of mature barrier cells^2–5^. Impaired epithelial renewal abates barrier function, allowing lumenal microbes to invade interstitial tissue, and promote disease. Thus, it is essential that we understand how enteric pathogens impact epithelial proliferation.

*Vibrio cholerae* (*Vc*) is a non-invasive, toxin-producing bacterium widely found in aquatic environments. Intestinal infection with *Vc* causes symptoms that range from mild gastrointestinal distress to the potentially fatal diarrheal disease, cholera. There are an estimated three million cholera cases with one hundred thousand deaths annually in endemic regions^6^. While antibiotics and vaccines protect from *Vc*, antibiotic-resistant strains are emerging, and current vaccines provide moderate, and time-limited protection^7^. We feel that a complete molecular profile of the gut response to *Vc* may identify pharmacologically relevant therapeutic targets with the potential to diminish or protect from infection.

The vinegar fly, *Drosophila melanogaster,* is a powerful system to study *Vc* pathogenesis^8^. Insects are natural *Vc* reservoirs and contribute to pathogen dissemination in endemic areas^9, 10^. Like the vertebrate small intestine, the fly midgut epithelium exists as a monolayer maintained by multipotent ISCs that typically proliferate to generate a daughter stem cell and transient cell types that mature into absorptive enterocytes or secretory lineages^11–16^. Fly ISCs predominantly generate a committed enterocyte precursor known as the enteroblast (EB), and EB-ISC pairs are defined as the progenitor compartment. Flies infected with *Vc* develop cholera-like symptoms that include epithelial breaches and watery diarrhea laden with live pathogen^17–19^. Like vertebrates, loss of cholera toxin does not completely abolish *Vc* virulence in flies, indicating cholera toxin-independent contributions to pathogenesis^17,18,20^. Flies have provided particularly valuable insights into the actions of *Vc* virulence factors. For example, polysaccharide-dependent biofilm formation allows *Vc* to colonize and persist in the fly gut^21^. The CtxA component of cholera toxin disrupts enterocyte junctions, leading to leakage, weight loss and reduced viability^22^. Furthermore, infection with *Vc* activates the fly Immune Deficiency (IMD) pathway^18,23^, a key mediator of gut antibacterial defenses with extensive similarities to the vertebrate Tumor Necrosis Factor Receptor (TNFR) response^24,25^. *Vc*-dependent activation of IMD contributes to pathogenesis by blocking ISC proliferation and promoting delamination of damaged epithelial cells, effectively preventing barrier repair in infected hosts^18,19,23^.

At present, it is unclear how IMD blocks proliferation, however a recent study showed that the *Vc* Type VI Secretion System (T6SS) is essential for arresting ISC division^26^. The T6SS is a syringe-like contractile apparatus that injects toxic effectors into susceptible prey, rapidly killing the target^27–30^. *Vc* uses the T6SS to eliminate competitors and enhance gut colonization^31–33^. T6SS-dependent killing of commensal bacteria contributes to *Vc* virulence, as inactivation of the T6SS or removal of gut commensals restores epithelial renewal, attenuates intestinal damage, and extends host lifespan after infection^19,26,33,34^. However, the underlying molecular mechanism of T6SS-dependent arrest of epithelial repair remains unclear.

Previously, we found that *Vc* enhances transcriptional expression of multiple Bone Morphogenetic Protein (BMP) pathway components in the fly progenitor compartment in a T6SS-dependent manner^26^. We consider this discovery noteworthy, as BMP members of the Transforming Growth Factor-β (TGF-β) cytokine family regulate intestinal homeostasis in flies and vertebrates ^35–46^. In both systems, BMPs act in a paracrine manner to inhibit progenitor proliferation and promote intestinal epithelial cell differentiation. Loss of BMP signaling in fly EBs leads to hyperproliferation of ISCs under normal conditions and upon infection with the entomopathogen *Erwinia carotovora carotovora 15* (*Ecc15*)^40^.

In this study, we used *Drosophila* to determine if the *Vc* T6SS engages host BMP signas to block intestinal epithelial renewal. We discovered that the T6SS is essential for BMP activation in progenitors, with indispensable contributions from commensal microbes and host IMD activity. Mechanistically, we showed that *Vc* specifically activates BMP in EBs, leading to a non-autonomous arrest of ISC proliferation. Together, our work resolves the mechanism by which *Vc* interacts with gut commensals and host innate defenses to prevent repair of the intestinal epithelial barrier. Notably, we extended this study to demonstrate that the T6SS also activates BMP and blocks ISC proliferation in a zebrafish infection model, suggesting an evolutionarily conserved impact of the T6SS on regeneration of infected intestines.

## RESULTS

### The T6SS Induces BMP Activation in *Drosophila* Intestinal Progenitors

In contrast to most enteric challenges, *Vc* causes widespread Intestinal Epithelial Cell (IEC) destruction without compensatory expansion of the progenitor pool^23,26^. At present, we do not understand how *Vc* blocks such an essential epithelial repair response. Recently, we showed that *Vc* enhances progenitor cell expression of multiple BMP pathway components in a T6SS-dependent fashion, including the ligand *decapentaplegic*, the type-I receptor *thickveins* (*tkv*), and the transcriptional target *spalt*^26^. As BMP is an evolutionarily conserved tumor suppressor that arrests IEC proliferation in flies and vertebrates, we tested the hypothesis that *Vc* prevents intestinal epithelial repair by activating BMP in progenitors.

In preliminary experiments, we used immunofluorescence-based imaging to determine the rostro-caudal distribution of GFP-expressing *Vc* in infected adult female midguts. We defined four levels of colonization that ranged from absent (level I) to extensive, biofilm-like accumulations (level IV, Figure S1A). We observed *Vc* distribution throughout the intestine, with prominent accumulations in the posterior midgut (Figure S1B), including IEC-associated *Vc* populations that had breached the peritrophic matrix (Figure S1C). As the posterior midgut is a region of high tissue turnover during infection, and *Vc* accumulates to the greatest extent in the posterior midgut, we elected to characterize relationships between *Vc,* BMP, and posterior midgut renewal in greater detail.

To determine if the T6SS activates BMP in host IECs, we quantified MAD phosphorylation (pMAD) in guts of uninfected *esg^ts^/+* flies, alongside *esg^ts^/+* flies that we challenged with the El Tor C6706 *Vc* strain, or the isogenic T6SS-deficient *vasK* deletion mutant (C6706Δ*vasK*)^47^. MAD phosphorylation provides a direct measure of BMP activation, and *esg^ts^/+* transgenic flies allow us to identify progenitors (small, GFP-positive cells), enteroendocrine cells (Prospero (Pros)-positive and GFP-negative), and enterocytes (large, GFP and Pros-negative cells), effectively allowing us to quantify lineage-specific BMP activation (Figure 1A). Regardless of infection status, we observed BMP activation 70-100% of Pros+ cells, indicating *Vc*-independent activation of BMP in the enteroendocrine lineage (Figures 1A and S2). In contrast, we detected substantial, T6SS-specific impacts on BMP activation within the progenitor compartment. Uninfected *esg^ts^/+* guts contained evenly spaced IECs with variable, but generally low levels of BMP activation throughout the epithelium and in progenitors (Figures 1A-1C). In agreement with earlier reports^23,26^, infection with wild-type *Vc* did not prompt expansion of progenitor cell numbers, indicating a failure of epithelial renewal (Figures 1A and S3). In contrast to uninfected guts, the average percentage of pMAD+ progenitors almost doubled in C6706-infected guts (Figures 1A-1C). Consistent with our previous study^26^, challenges with C6706Δ*vasK* stimulated progenitor growth (Figures 1A and S3). However, we did not observe effects of C6706Δ*vasK* infection on MAD phosphorylation in gut epithelial cells or the progenitor compartment relative to uninfected controls (Figures 1A-C), suggesting that T6SS-deficient *Vc* fails to modify the host BMP pathway. Combined with our earlier transcriptional profiles^26^, our data indicate a T6SS-specific activation of BMP in the progenitor compartment with an accompanying failure to expand progenitor numbers during infection.

**Figure 1.**
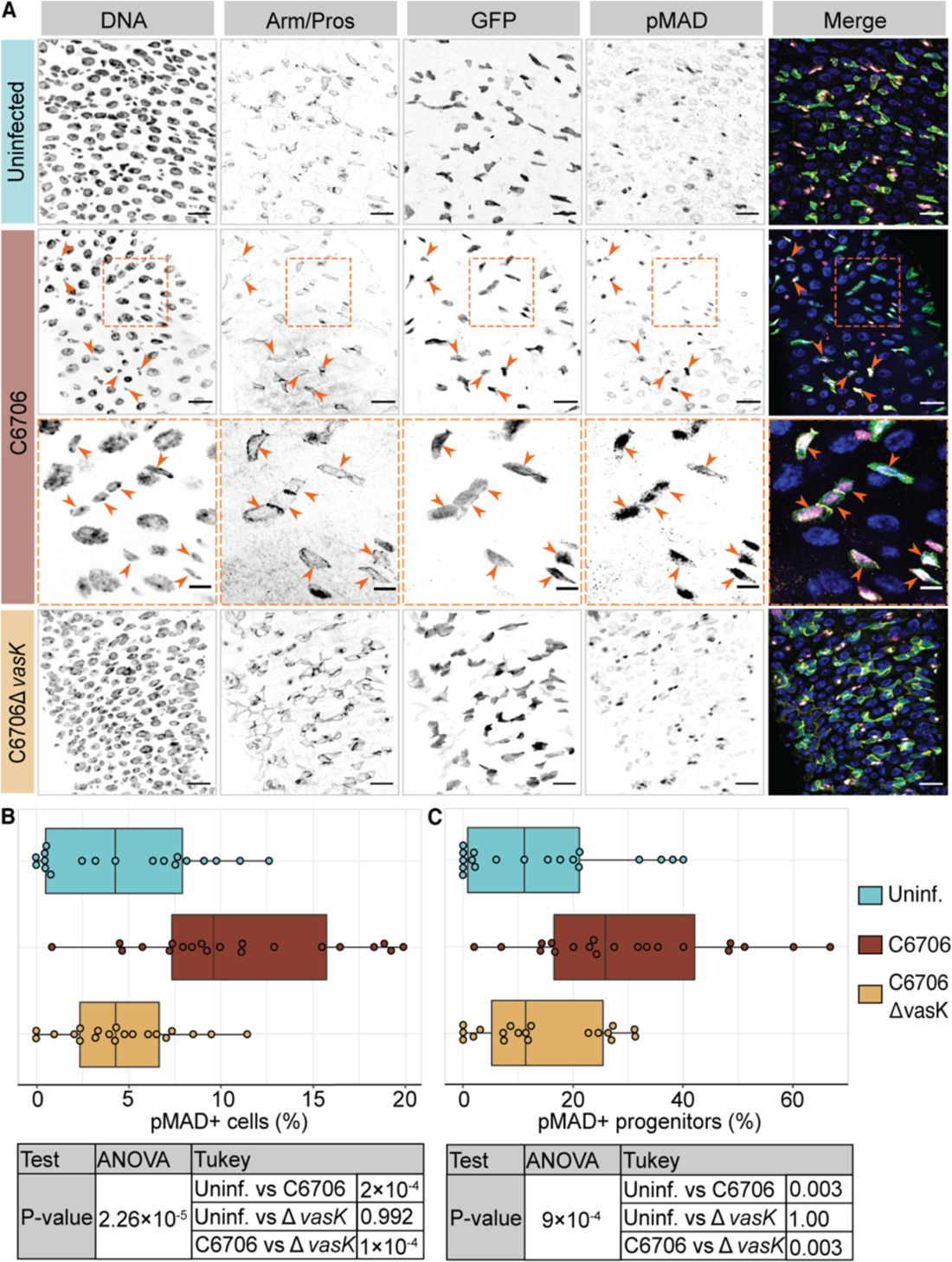
*Vc* activates BMP in intestinal progenitor cells in a T6SS-dependent manner. **(A)** Posterior midguts of *esg^ts^/+* adult fly uninfected, infected with C6706 or C6706Δ*vasK* for 24 h. Hoechst labels DNA (blue), Armadillo and Prospero (Arm/Pros) label cell borders and enteroendocrine cells, respectively (yellow), GFP labels progenitors (green), and pMAD labels cells with BMP activation (magenta). Arrows and dashed boxes = pMAD+ GFP+ progenitors. Scale bars in dashed boxes = 5 μm. Scale bars in images = 15 μm. **(B and C)** Percentage of **(B)** all cells or **(C)** progenitors that are pMAD+. Each dot represents a measurement from a single fly gut. P values are calculated using the significance tests indicated in the tables.

### *Vc-*Mediated Activation of BMP Requires Commensal Bacteria and Host Innate Defenses

As T6SS-dependent pathogenesis involves interactions between *Vc* and commensal bacteria in flies and mice^19,33^, we asked if commensals are also required for T6SS-responsive BMP activation. In this instance, we quantified infection-dependent BMP activation in germ-free (GF) *esg^ts^/+* flies relative to conventionally reared (CR) counterparts. On average, we observed a three-fold increase in pMAD+ progenitor numbers of infected CR flies compared to uninfected controls (Figures 2B and 2C), with a parallel failure to increase progenitor numbers (Figures 2B and 2D), further supporting the argument that *Vc* activates BMP while arresting cell proliferation. However, in the absence of gut-associated bacteria, infection increased progenitor numbers by an average of 25% (Figures 2A and 2D), without activation of BMP (Figures 2A and 2C), establishing a requirement for gut commensals to support *Vc*-dependent activation of BMP and arrest epithelial repair.

**Figure 2.**
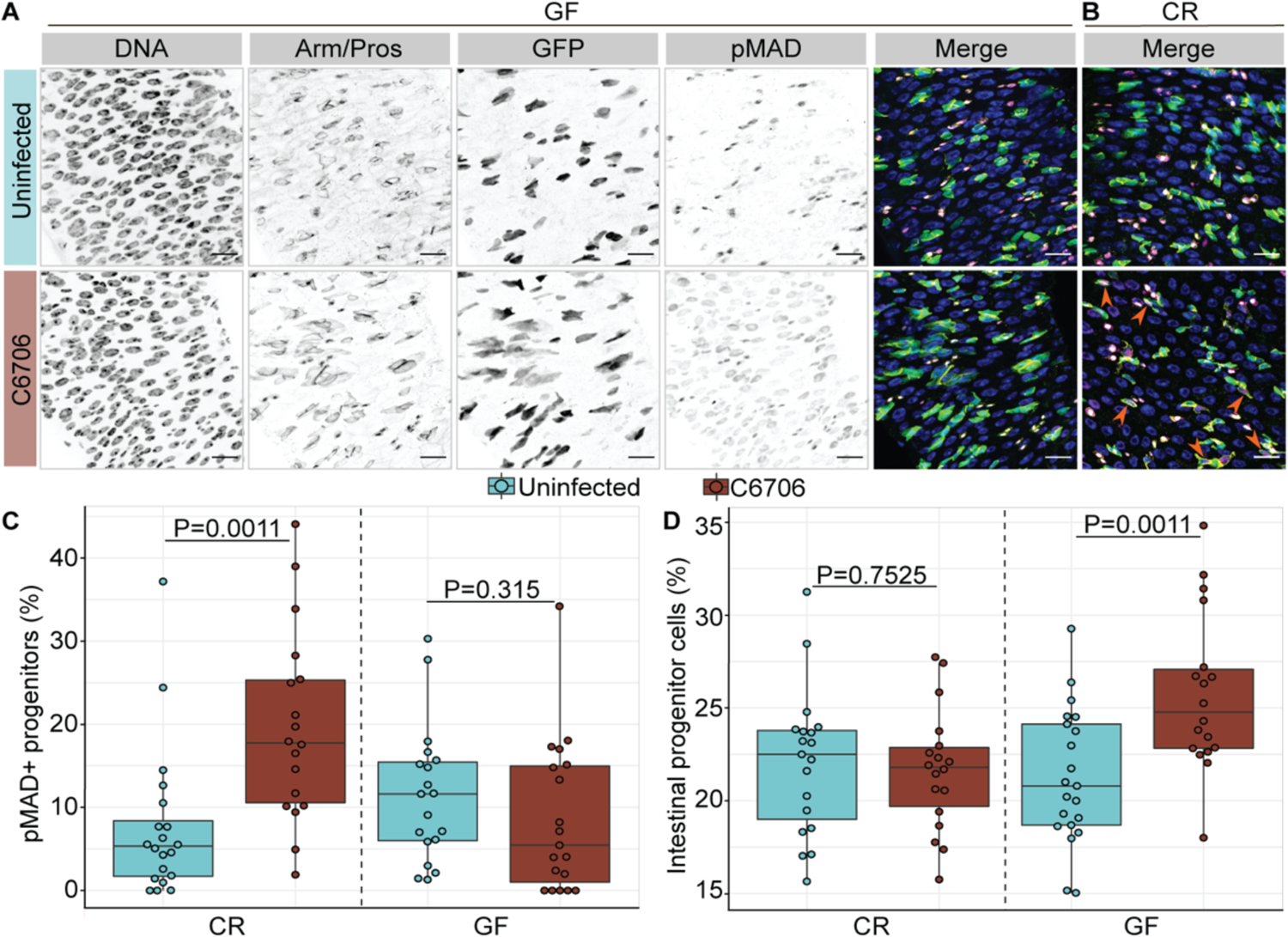
Progenitor-specific activation of BMP signaling requires *Vc*-commensal interactions. **(A and B)** Posterior midguts of **(A)** germ-free (GF) or **(B)** conventionally reared (CR) *esg^ts^/+* adult flies uninfected or infected with C6706 for 24 h. Hoechst labels DNA (blue), Armadillo and Prospero (Arm/Pros) label cell borders and enteroendocrine cells, respectively (yellow), GFP labels progenitors (green), and pMAD labels cells with BMP activation (magenta). Scale bars = 15 μm. **(C)** Percentage of GFP+ progenitors that are pMAD+ in GF and CR *esg^ts^/+* flies. **(D)** Proportion of GFP+ progenitors in all intestinal epithelial cells in GF and CR *esg^ts^/+* flies. Each dot represents a measurement from a single fly gut. P values are calculated using unpaired Student t-tests.

As the TNFR-like Immune Deficiency (IMD) pathway orchestrates gut responses to commensal bacteria^3,48^, and IMD prevents ISC proliferation upon *Vc* infection^19,23^, we asked if T6SS-commensal interactions activate BMP via IMD. To probe links between *Vc*, IMD, and BMP, we compared progenitor-specific BMP activation in wild-type and *imd* null mutant flies that we challenged with *Vc*. Our *imd* mutant line and isogenic wildtype control do not express GFP under control of the *esg* promoter. Therefore, we defined progenitors as small, Armadillo-enriched, Prospero-negative cells based on established cellular characteristics. In agreement with Figures 1 and 2, we found that *Vc* increased pMAD+ progenitor numbers in wildtype host midguts (Figures 3A and 3C). However, we also discovered that *Vc* failed to activate BMP in midguts of IMD-deficient flies, suggesting a role for IMD in BMP activation (Figures 3B and 3C). To identify the epithelial cell type where IMD acts to stimulate a host BMP response, we monitored BMP activation levels in intestines of flies with cell type-specific depletion of the IMD-responsive NF-κB family transcription factor *relish* (*rel*). Inactivation of *rel* in progenitors (*MyoIA^ts^>rel^RNAi^*) did not have appreciable consequences for infection-dependent MAD phosphorylation (Figure 3D), suggesting that *Vc*-dependent BMP activation does not require an IMD-Relish response in progenitors. In contrast, we did not observe infection-dependent increases in MAD phosphorylation upon enterocyte-specific inactivation of *rel* (*MyoIA^ts^>rel^RNAi^*, Figure 3E), indicating enterocyte-specific requirements for IMD in BMP activation during infection. As a whole, our data suggest that T6SS-dependent activation of progenitor-specific BMP requires pathogen-commensal interactions and IMD activity in enterocytes.

**Figure 3.**
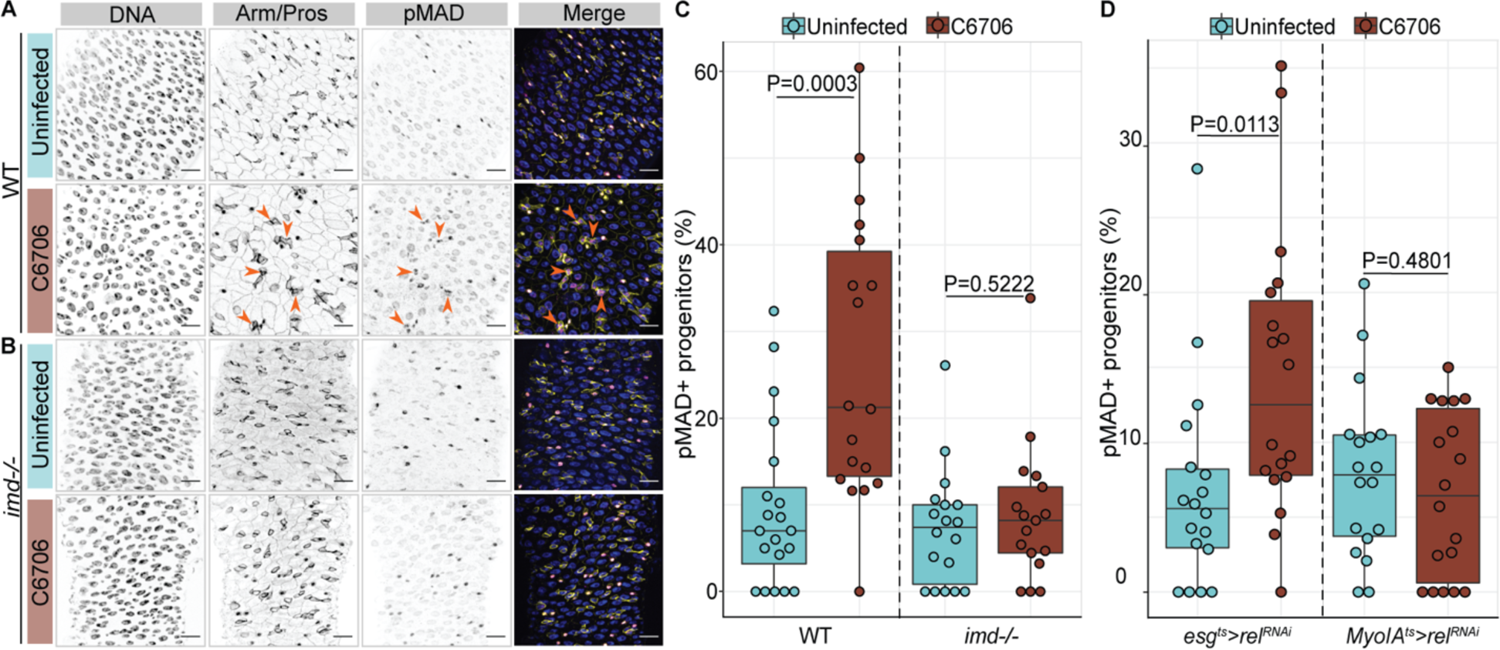
*Vc-*responsive activation of BMP requires IMD activity. **(A and B)** Posterior midguts of **(A)** wild-type (WT) or **(B)** *imd* null mutant *(imd-/-*) adult flies uninfected or infected with C6706 for 24 h. Hoechst labels DNA (blue), Armadillo and Prospero (Arm/Pros) label cell borders and enteroendocrine cells, respectively (yellow), and pMAD labels cells with BMP activation (magenta). Arrows = pMAD+ progenitor cells. Scale bars = 15 μm. **(C)** Percentage of progenitors that are pMAD+ in WT and *imd-/-* flies. **(D)** Percentage of pMAD+ progenitors in *esg^ts^>rel^RNAi^* and *Myo^ts^>rel^RNAi^* flies. Each dot represents a measurement from a single fly gut. P values are calculated using unpaired Student t-tests.

### *Vc*-Dependent Activation of BMP is Essential to Arrest Epithelial Repair in Infected Flies

Thus far, our data are consistent with a link between BMP activation by the T6SS and an arrest of progenitor cell expansion. However, we lack phenotypic data that specifically determines if BMP activation is essential for T6SS-dependent progenitor growth arrest. As T6SS-deficient *Vc* fail to activate BMP or arrest progenitor expansion (e.g., Figure 1C), we first tested if progenitor-specific activation of BMP blocks ISC proliferation in flies challenged with T6SS-deficient *Vc*. Specifically, we compared progenitor cell expansion in control *esg^ts^/+* flies and in flies that expressed a constitutively active variant of the type-I BMP receptor *tkv* (*esg^ts^>tkv^CA^*) in progenitors. We observed an approximately 25% expansion of GFP+ progenitor cells in infected *esg^ts^/+* guts, confirming activation of the host repair response (Figures 4A and 4C). In contrast, T6SS-deficient *Vc* failed to stimulate compensatory progenitor growth in *esg^ts^>tkv^CA^* guts (Figure 4B-C), demonstrating that progenitor-specific BMP activation is sufficient to arrest proliferation in an infected host.

**Figure 4.**
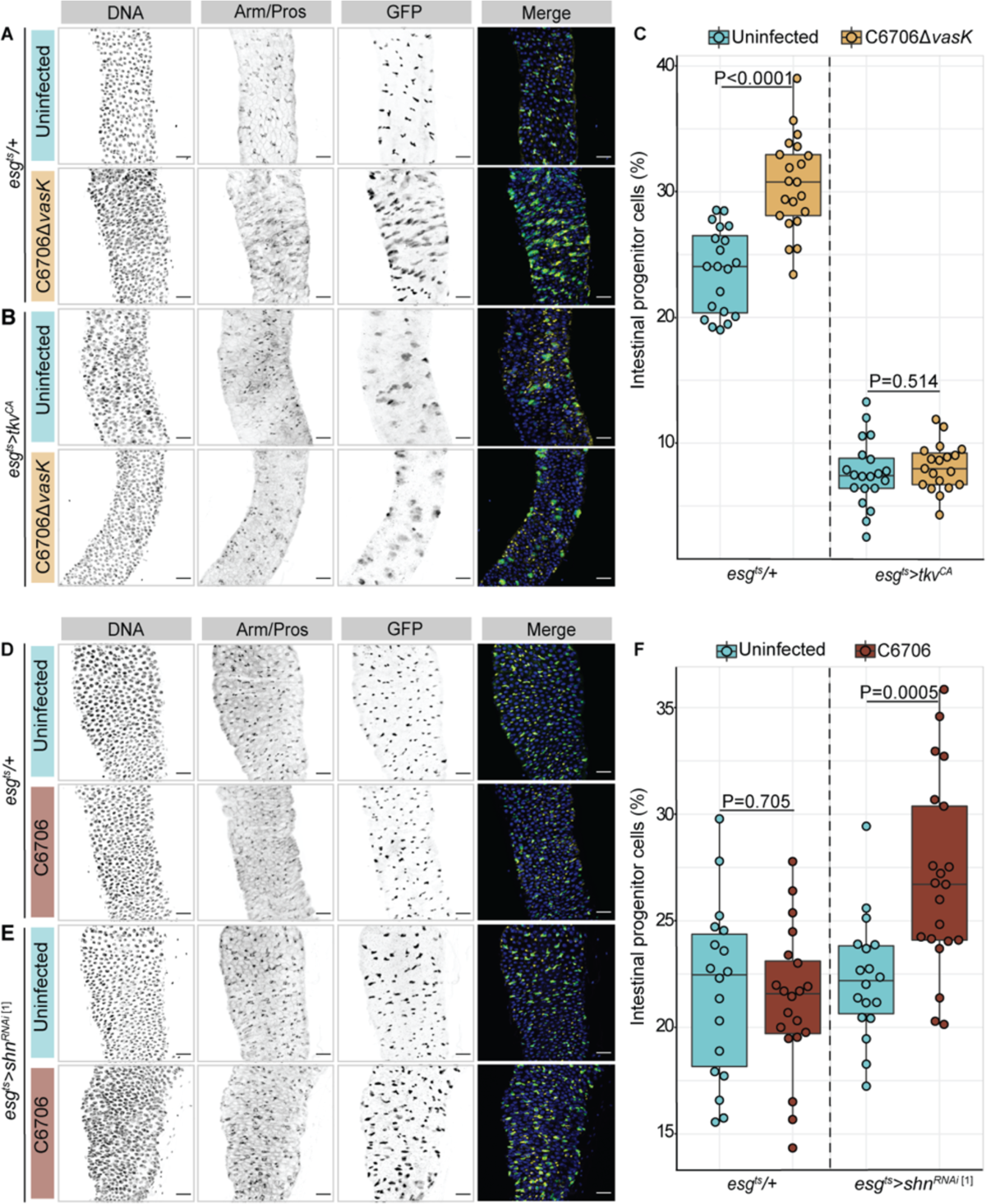
BMP regulates epithelial repair after *Vc* infection. **(A and B)** Posterior midguts from **(A)** wide-type (*esg^ts^/+*) or **(B)** progenitor-specific BMP activation (*esg^ts^>tkv^CA^*) adult flies uninfected or infected with C6706Δ*vasK* for 36h. DNA labeled with Hoechst (blue), GFP marks progenitors (green), Armadillo and Prospero (Arm/Pros) label cell borders and enteroendocrine cells, respectively (yellow). Scale bar = 25 μm. **(C)** Proportion of intestinal epithelial cells that are GFP+ progenitors in *esg^ts^/+* and *esg^ts^>tkv^CA^* flies. **(D and E)** Images of **(D)** wide-type (*esg^ts^/+*) or **(E)** progenitor-specific BMP inhibition (*esg^ts^>shn^RNAi^* <u>^[1]^</u>) adult flies uninfected or infected with C6706 for 36h. DNA labeled with Hoechst (blue), GFP marks progenitors (green), Armadillo and Prospero (Arm/Pros) label cell borders and enteroendocrine cells, respectively (yellow). Scale bar = 25 μm. **(F)** Proportion of cells that are GFP+ progenitors in *esg^ts^/+* and *esg^ts^>shn^RNAi^* flies. Each dot represents a measurement from a single fly gut. P values are calculated using unpaired Student t-tests.

In addition to preventing C6706Δ*vasK*-driven progenitor cell expansion, activation of the BMP pathway had visible effects on the intestinal epithelium of uninfected flies (Figure 4B), including a substantial decline in the number of intestinal progenitors (Figure 4C). Thus, while interesting, effects of *esg^ts^>tkv^CA^* on host responses to T6SS-deficient *Vc* do not adequately establish a requirement for BMP to arrest progenitor renewal after infection. To directly test if BMP modifies host responses to *Vc*, we specifically ablated BMP signaling in the progenitor compartment by knocking down the transcription factor *shn* (*esg^ts^>shn^RNAi^*) and monitored progenitor numbers in flies that we challenged with wild-type *Vc*. As anticipated, *Vc* did not stimulate compensatory progenitor expansion or cell proliferation in infected *esg^ts^/+* flies (Figures 4D, 4F, S4A). However, loss of BMP activity in progenitors effectively ablated *Vc*-mediated inhibition of progenitor growth. Instead, we found that infected *esg^ts^>shn^RNAi^* flies contained more progenitors and more PH3+ proliferating cells than uninfected counterparts, similar to what we saw in wild-type guts upon challenge with T6SS-deficient *Vc* (Figures 4D-F and S4A-G). We saw similar consequences of *shn* inactivation with two alternative *shn^RNAi^* lines, confirming that BMP is necessary in the progenitor compartment to arrest epithelial renewal upon *Vc* infection.

### BMP-dependent arrest of proliferation requires a non-autonomous signal from EBs to ISCs

As progenitor cells consist of self-renewing ISCs and transient EBs, we next tested the involvement of each cell type in BMP-dependent arrest of epithelial repair. To determine if *Vc* activates BMP in a specific progenitor cell type, we challenged EB-specific driver line *Su(H)^ts^/+* flies with C6706 and quantified the number of pMAD+ ISCs (Dl+, GFP-) and pMAD+ EBs (Dl-, GFP+) in posterior midguts. In both infected and uninfected guts, there were similar and relatively low amounts of ISCs with BMP activity (Figures 5A and 5B). In contrast, infection with C6706 stimulated variable, but significant, activation of BMP in EBs, with an average threefold increase in the percentages of pMAD+ EBs (Figures 5A and 5B), indicating that *Vc* infection preferentially activates BMP in EBs. To test impacts of EB-specific BMP activation on epithelial renewal during infection, we knocked down *shn* exclusively in EBs (*Su(H)^ts^>shn^RNAi^*) and quantified progenitors after infection. We discovered that, like progenitor-wide knockdown, EB-specific loss of *shn* ablated *Vc*-responsive arrest of progenitor growth (Figure 5C). Moreover, we observed an increase in the proportion of ISCs, with no significant changes in EB numbers in infected guts, suggesting that BMP in EBs regulates ISC proliferation upon challenge with *Vc* (Figures 5D and 5E).

**Figure 5.**
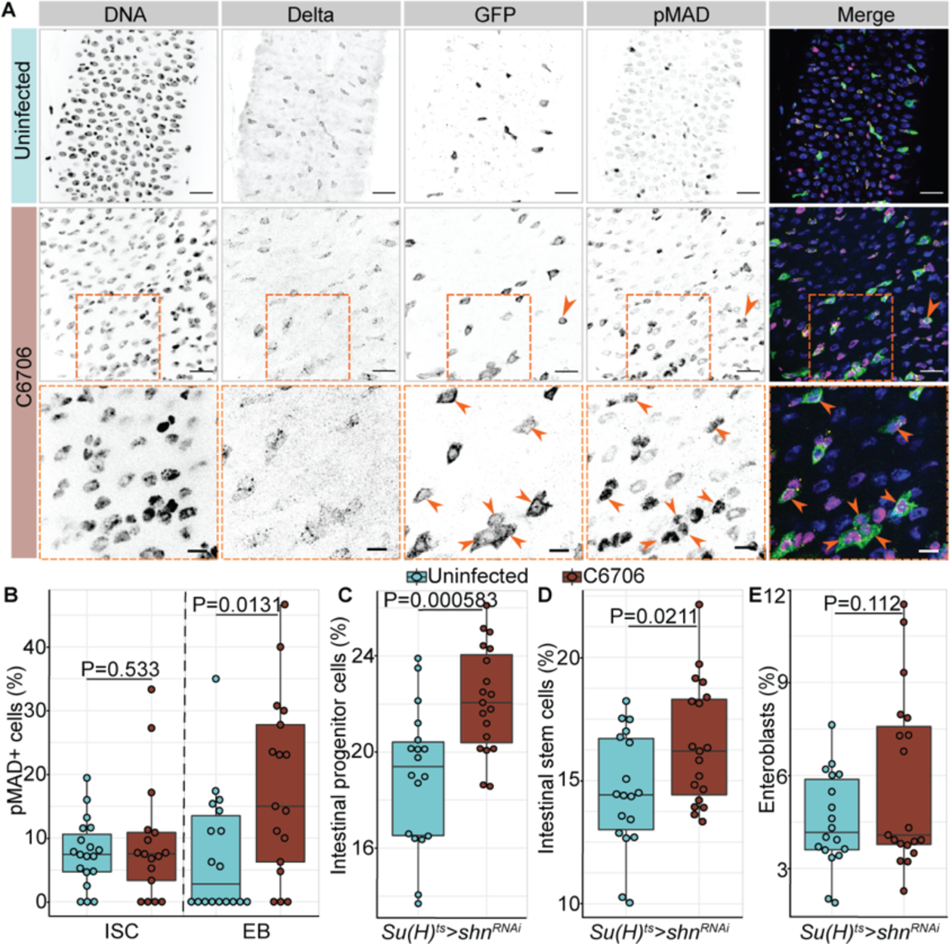
*Vc*-responsive BMP activation in EBs arrests intestinal stem cell growth non-cell autonomously. **(A)** Posterior midguts *Su(H)^ts^/+* adult flies of uninfected or infected with C6706 for 36h. DNA marked by Hoechst (blue), Delta+ intestinal stem cells (yellow), Su(H)+ EBs marked by GFP (green), and pMAD (magenta) to monitor BMP activation. Arrows and dashed box = pMAD+ GFP+ EBs. Scale bars in dashed boxes = 5 μm. Scale bars in images = 15 μm. **(B)** Proportion of intestinal stem cells or EBs that are pMAD+ in *Su(H)^ts^/+* intestines. **(C-E)** Proportions of all cells that are **(C)** progenitors, **(D)** intestinal stem cells, or **(E)** EBs in the intestines with EB-specific BMP inactivation (*Su(H)^ts^>shn^RNAi^*). Each dot represents a measurement from a single fly gut. P values are calculated using Student t-tests.

### The T6SS suppresses cell proliferation and induces TGF-β/BMP activation in a vertebrate intestine

To expand our study, we then asked if the impact of the *Vc* T6SS on intestinal epithelial renewal applies to vertebrate hosts. Zebrafish *Danio rerio* are ideal vertebrate models to study *Vc* pathogenesis^49–51^. Fish intestines are naturally colonized by *Vibrio* species and develop cholera-like symptoms upon *Vc* infection^49,51–54^. Importantly, like the mammalian intestine, the zebrafish intestinal epithelium contains a complex community of secretory and absorptive lineages that interact with immune-regulatory myeloid and lymphoid cells^55–59^. To test whether the *Vc* T6SS impacts epithelial repair in fish, we challenged zebrafish larvae and adults with either wildtype *Vc* (C6706) or T6SS-deficient *Vc* (C6706Δ*vasK*) and measured cell proliferation. In larvae, uninfected intestines contained moderate amounts of EdU+ proliferating cells (Figures 6A and 6B). Infection with C6706Δ*vasK* promoted epithelial repair indicated by a significantly higher proliferation rate compared to uninfected guts (Figures 6A and 6B). In contrast, there was no significant difference in numbers of EdU+ cells between C6706-infected and uninfected guts. In parallel to our findings in larvae, we also discovered that the *Vc* T6SS disrupts IEC regeneration in adult zebrafish intestines. Both C6706 and C6706Δ*vasK* damaged the intestinal epithelium, characterized by disorganized IECs and lumenal shedding of damaged tissue (Figure 6C, representative instances of barrier damage or lumenal shedding highlighted with arrowheads). Damage in C6706Δ*vasK-* infected guts stimulated repair, as the numbers of PCNA+ proliferating cells nearly doubled compared to uninfected counterparts (Figures 6C-D). In contrast, we did not detect significant changes in cell proliferation rates after C6706 infection (Figures 6C-D). These data demonstrate that, like a fly host, *Vc* arrests epithelial proliferation in zebrafish intestines in a T6SS-dependent manner. Furthermore, we found that *the* T6SS induced TGF-β/BMP, as larvae infected with wildtype *Vc* contained significantly larger numbers of cells with the active, phosphorylated form of SMAD3 (pSMAD3) compared to uninfected and T6SS-deficient *Vc* infected counterparts (Figures 6E and 6F).

**Figure 6.**
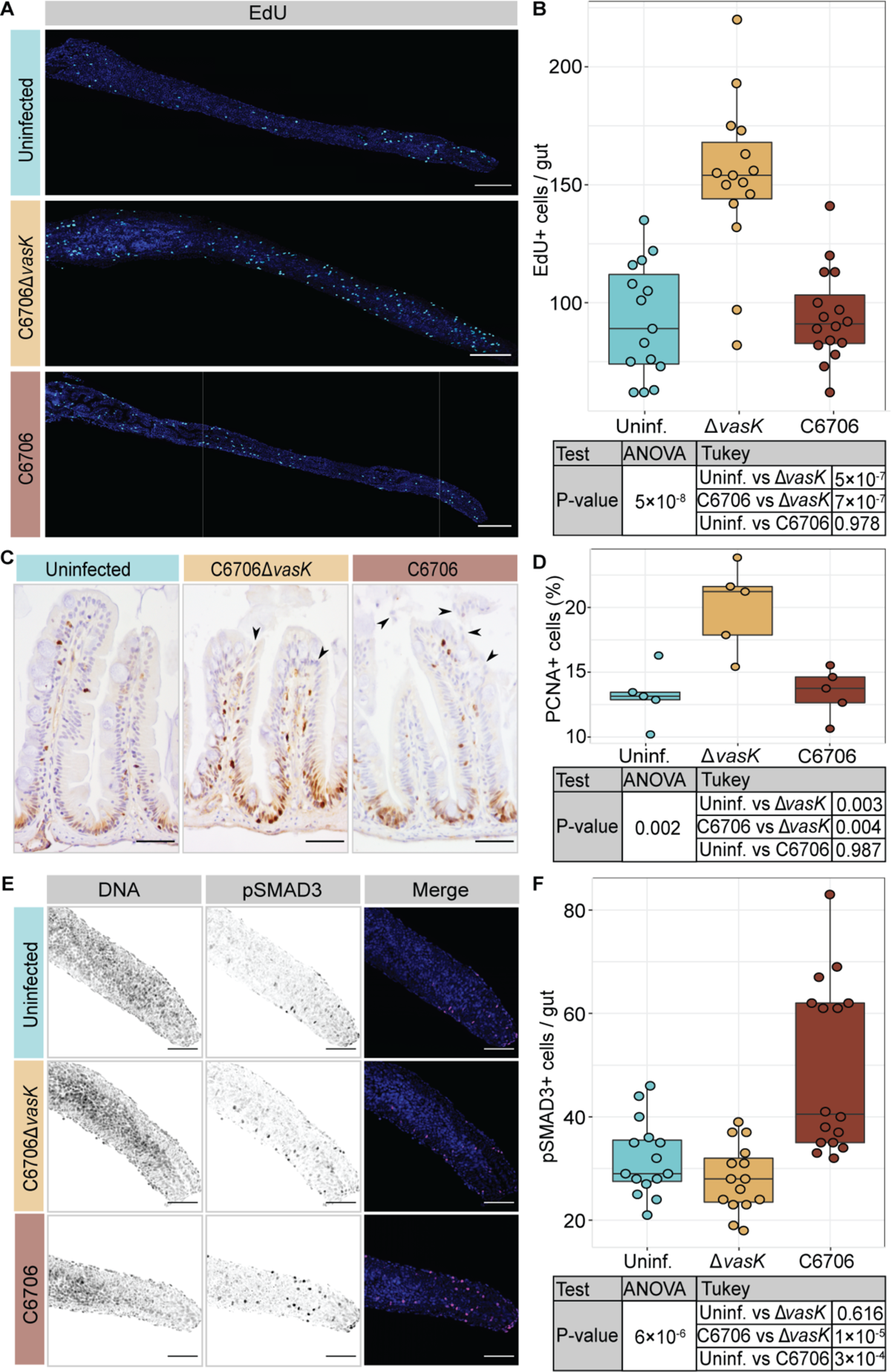
*Vc* T6SS induces BMP activation and limits cell proliferation in the zebrafish intestine. **(A)** Intestines of TL zebrafish larvae uninfected or *Vc*-infected for 24h with DNA stained by Hoechst in blue and EdU+ cells in cyan. Scale bars = 200 μm. **(B)** Quantification of EdU+ cells per gut. **(C)** Immunohistochemical images of sagittal posterior intestinal sections from adult TL zebrafish uninfected or *Vc-*infected for 24h stained for PCNA. Arrows indicate epithelial damages marked by disorganized nuclei and shedding of epithelial cells. Scale bars = 50 μm. **(D)** Percentage of intestinal epithelial cells that are PCNA+ in adult fish guts. **(E)** Posterior intestines from TL zebrafish larvae uninfected or *Vc*-infected for 24h with DNA marked by Hoechst in blue and pSMAD3 in magenta. Scale bars = 100 μm. **(F)** Quantification of pSMAD3+ cells per gut. Each dot represents a measurement from a single fish intestine. P values are calculated using the significance tests indicated in the tables.

As *Vc* blocks intestinal cell proliferation in a T6SS-dependent manner, we asked if pathogen-commensal interactions are necessary for *Vc-*responsive arrest of epithelial repair in a zebrafish host. To answer this question, we quantified proliferating cells in uninfected, or infected, germ-free (GF) intestinal epithelia relative to conventionally reared (CR) counterparts. Like flies, infection with C6706 failed to stimulate tissue repair in CR fish, as unchallenged and infected guts contained similar amounts of EdU+ cells (Figures 7A and 7C). In agreement with earlier reports that the microbiota stimulates epithelial proliferation in zebrafish intestines ^60,61^, we observed fewer EdU+ cells in uninfected GF fish than in uninfected CR controls (Figure 7B). However, in contrast to CR counterparts, C6706 infection induced a more than threefold increase in EdU+ cell numbers in GF guts relative to CR guts, indicating that *Vc* fails to block epithelial repair in the absence of gut commensals (Figures 7B and 7C). Although further studies are needed to determine whether TGF-β/BMP activity is needed for T6SS-responsive arrest of cell proliferation, our findings raise the possibility that *Vc* prevents repair of damaged intestinal epithelia through similar mechanisms in invertebrates and vertebrates.

**Figure 7.**
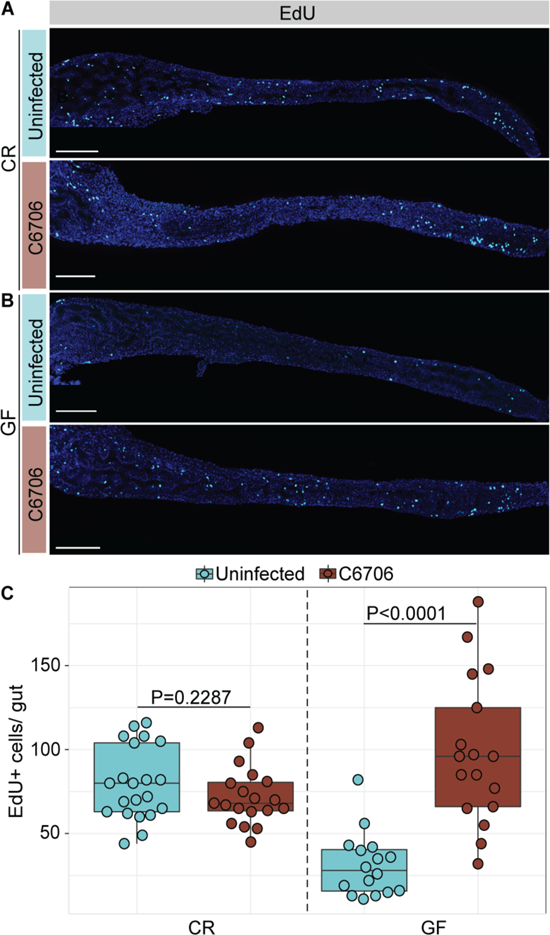
*Vc* T6SS-responsive arrest of cell proliferation requires commensals in the zebrafish intestine. **(A and B)** Intestines of **(A)** conventionally reared (CR) or **(B)** germ-free (GF) TL zebrafish larvae uninfected or infected with C6706 for 24 h. Hoechst labels DNA (blue) and EdU labels proliferating cells (cyan). Scale bars = 200 μm. **(C)** Quantification of EdU+ cells per gut. Each dot represents a measurement from a single larval intestine. P values are calculated using unpaired Student t-tests.

## DISCUSSION

Enteric infection normally accelerates cell proliferation and tissue repair via stress responses such as the epidermal growth factor receptor (EGFR) and Janus kinase-signal transducer and activator of transcription (JAK-STAT) pathways^3–5,62^. However, oral infection of *Drosophila* with *Vc* leads to T6SS-dependent arrest of intestinal proliferation^26^. It is unknown how *Vc*-commensal interactions block epithelial repair. We consider this an important question, as failure to renew the intestinal epithelium exposes peri-intestinal tissue to gut-resident microbes, greatly increasing the risk of systemic infection. In this study, we found that interactions between *Vc* and host commensals activate BMP specifically in intestinal progenitor cells through the TNFR-like IMD pathway. BMP activity in EBs arrests ISC proliferation non-autonomously, which is necessary to block epithelial repair. Our study reveals how pathogen-commensal interactions engage evolutionarily conserved immune and growth regulators to impact epithelial renewal and also raises several interesting aspects to be further elucidated.

How does IMD activate intestinal BMP during a *Vc* infection? Normally, IMD serves a protective role and is essential for surviving infections by pathogens like *Pseudomonas entomophila*^63–65^. In contrast, *Vc*-dependent activation of IMD arrests ISC proliferation and shortens the host lifespan^18,19,23^. We found that *Vc*-commensal interactions activate BMP in progenitors via IMD signaling in enterocytes. IMD is typically activated by bacterial diaminopimelic acid-type peptidoglycan (DAP-PGN)^66–71^, although alternate mechanisms also exist in the gut^72^. We do not know which commensal species are responsible for initiating an IMD-BMP response in the fly gut. Fly symbiont *Acetobacter sp.* contains DAP-PGN and *Acetobacter pasteurianus* is sensitive to *Vc* T6SS^19^. In addition, T6SS-dependent killing of *A. pasteurianus* accelerates host death and contributes to *Vc* pathogenesis^19^. Therefore, it is possible that microbial components generated by T6SS-*Acetobacter sp.* interactions initiate IMD response in the gut. However, interactions between *Vc* T6SS and *A. pasteurianus* alone are insufficient to block intestinal epithelial repair^26^, indicating that other commensal species are involved in *Vc*-responsive activation of an IMD-BMP axis to impact tissue renewal. As gut microbes are essential for BMP-dependent arrest of epithelial proliferation, we believe that variability within the host microbiome may explain the inter-fly variability in MAD phosphorylation, and progenitor pool expansion observed in many of our experiments. The fly gut microbiome is constantly replenished through feeding^73^, and bacterial acquisition by the host is stochastic^74^, and variable between individual hosts^75^. Given the ease of reconstituting gut microbiota in *Drosophila*, further studies in gnotobiotic flies would be helpful to identify the commensal species involved in *Vc*-responsive activation of an IMD-BMP response.

How does BMP regulate ISC proliferation? Mechanistically, our data suggest a non-autonomous role of BMP signaling in regulating epithelial renewal during infection. We found that *Vc*-responsive BMP activation in EBs blocks proliferation of neighboring ISCs. Our findings match an earlier report that EB-specific BMP inactivation stimulates ISC hyperproliferation upon challenges with *Ecc15* or in the absence of infection^40^. Currently, it is unclear how EB-specific BMP activity regulates ISC proliferation. Typically, epithelial damage during enteric infections enhance JAK-STAT and EGFR signals in ISCs, leading to an elevated proliferation rate^3–5,62^. A recent study suggested a functional link between BMP and stress responses in regulating intestinal epithelial renewal. Specifically, loss of BMP signals activated of JAK-STAT and EGFR signaling in fly midguts, which fueled stem cell division and eventually disrupted epithelial homeostasis^39^. Our previous transcriptional data showed that wildtype *Vc* did not initiate JAK-STAT and EGFR activity in midgut progenitors, whereas T6SS-deficient *Vc* successfully initiated both responses^26^. Thus, we speculate that BMP activity in EBs blocks epithelial proliferation by antagonizing JAK-STAT or EGFR responses in neighboring ISCs upon *Vc* infection. Further investigations are required to elucidate the mechanisms by which EB-specific BMP non-autonomously regulates stem cell growth during infections.

How does BMP regulate epithelial homeostasis during enteric infections? We found that T6SS-responsive activation of BMP blocks ISC proliferation, leading to an arrest of epithelial repair. Notably, our findings are not restricted to flies. *Vc* T6SS induces TGF-β/BMP activity in zebrafish intestines. Moreover, infection with *Vc* suppresses cell proliferation in a manner that requires a T6SS and the presence of commensal microbes. It remains to be seen whether BMP signaling is necessary to block epithelial proliferation in the fish gut and whether the T6SS-responsive arrest of epithelial repair requires microbial interactions. Unlike fly guts which rely on germline-encoded innate defenses, zebrafish intestines also possess lymphocyte-based adaptive defenses that impact host responses to symbiotic and pathogenic microbes^76–82^. Recent studies reveal an immune-modulatory role of BMP in the vertebrate intestine^83–85^. In both human and mouse colons, epithelial-specific BMP activity limits expression of pro-inflammatory genes including a number of chemokines and cytokines and suppresses infiltration of macrophages and neutrophils^84,85^. Loss of the BMP pathway transcription factor SMAD4 in epithelial cells promotes inflammation-driven carcinogenesis in mouse colon^84,86^. Based on these findings and advantages of the zebrafish model, future studies could be conducted in fish to determine whether epithelial BMP signaling synergizes with innate and adaptive defenses to regulate the intestinal epithelial barrier during *Vc* infection. In addition to controlling cell proliferation and immune responses, intestinal BMP also regulates differentiation of immature intestinal epithelial cells in flies and vertebrates^37,44,87,88^. Therefore, it is possible that *Vc*-responsive BMP activation also impacts regeneration of a mature intestinal epithelium upon infection. Further investigations are needed to precisely determine roles of epithelial-specific BMP activity during enteric infections and how gut symbiotic and pathogenic microbes affect host responses.

In summary, this study demonstrates that *Vc*-commensal interactions modify host immune and growth regulatory pathways to disrupt intestinal epithelial repair. Using a *Drosophila* model, we uncovered effects of BMP activity and innate defenses on epithelial renewal upon infection with *Vc*. Given the importance of BMP and TNFR-like signaling in regulating intestinal homeostasis, we consider this work relevant to understand how evolutionarily conserved pathways orchestrate host responses to a global health threat.

## ACKNOWLEDGEMENTS

We are grateful to Bloomington Drosophila Stock Center, Dr. Bruce Edgar, Dr. Bruno Lemaitre, and Dr. Minjeong Shin for providing fly lines. *V. cholerae* C6706 was provided by Dr. John Mekalanos. We would also like to thank Science Animal Support Services and the Alberta Health Sciences Animal Laboratory Services for their excellent care of the zebrafish aquatic facilities. We acknowledge microscopy support from Dr. Steven Ogg, Gregory Plummer, and Kiera Smith at the Faculty of Medicine and Dentistry Cell Imaging Core. We acknowledge TEM support from Sara Amidian and Dr. Xuejun Sun at the Department of Oncology Cell Imaging Facility. We acknowledge the immunohistochemistry support provided by Dr. Kacie Norton at the Department of Biological Sciences Advanced Microscopy Facility and help from Mckenna Eklund. Thanks to Dr. Robert Ingham, Lena Jones, Mckenna Eklund, Aurélia Joly and Ralph Derrick De Leon for critical reading of the manuscript. This work was supported by a grant from the Canadian Institutes of Health Research to E.F. (MOP77746). X.X. has funding support through Alberta Graduate Excellence Scholarship and Li Ka Shing Institute of Virology Graduate Studies Entrance Award.

## MATERIALS AND METHODS

### Fly and zebrafish husbandry

Fly stocks and crosses were maintained at 18°C on standard cornmeal medium (Nutri-Fly Bloomington formulation https://bdsc-indianaedu.login.ezproxy.library.ualberta.ca/information/recipes/bloomfood. html). All experimental flies were virgin females kept at 18°C during collection and shifted to 29°C with a 12 hour/12 hour light/dark cycle to induce expression of transgenes of interest. Fly lines used in this study include *imd^EY08573^ (imd-/-)*^89^, *w;esg-GAL4,tubGAL80^ts^,UAS-GFP (esg^ts^)*^11^, *w;Su(H)GBE-GAL4,UAS-GFP;ubi-GAL80^ts^ (Su(H)^ts^)*^90^, *UAS-shn^RNAi^* ^[1]^ (VDRC #KK-105643), *UAS-shn^RNAi^* ^[2]^ (Bloomington #34689), *UAS-shn^RNAi^* ^[3]^ (Bloomington #82982), *UAS-tkv^CA^* (Bloomington #36536), *40D-UAS* (control for VDRC KK lines, VDRC #60101). We used *w^11^*^18^ as a wild-type control to set up crosses with *esg^ts^* and *Su(H)^ts^* lines. The *imd* line was backcrossed into our wild-type *w^1118^* background for eight generations prior to use. As *white* mutants showed impaired biological functions including stress response in flies^91^, we generated a wild-type control in Figure 3 for *imd-/-* by introducing the *w+* allele from *Oregon R* line into our *w^11^*^18^ via backcrossing for eight generations.

Zebrafish were raised and maintained following protocols approved by the Animal Care & Use Committees, Biosciences and Health Sciences at the University of Alberta, operating under the guidelines of the Canadian Council of Animal Care. Adult TL strain zebrafish were reared at the University of Alberta fish facility at 29°C under a 14-hour/10-hour light/dark cycle using standard zebrafish husbandry protocols. Zebrafish were fasted for 22 hours prior to infection. For larval analysis, TL zebrafish embryos were collected from breeding tanks and transferred to glass petri dishes with embryo media (15 mM NaCl, 1 mM CaCl_2_•2H_2_O, 1 mM MgSO4•7H_2_O, 0.7 mM NaHCO_3_, 0.5 mM KCl, 0.15 mM KH_2_PO_4_ and 0.05 mM Na_2_HPO_4_ in MiliQ water). Embryos were raised at 29°C under a 14-hour/10-hour light/dark cycle. 6 day-post-fertilization larvae were used for *Vc* infection.

### Generation of germ-free flies

To generate germ-free (GF) flies, freshly eclosed female *esg^ts^/+* flies were raised on autoclaved food with antibiotics (100µg/mL Ampicillin, 100µg/mL Metronidazole, 50µg/mL Vancomycin dissolved in 50% ethanol, and 100µg/mL Neomycin dissolved in water) for 5 days at 25°C. To ensure sterility, two flies from each vial were homogenized in MRS broth and plated on MRS agar plates. Flies were considered GF if no visible colonies formed. Conventionally reared (CR) flies were fed autoclaved food without antibiotics for 5 days at 25°C. Both GF and CR flies were then transferred onto autoclaved food without antibiotics for 7 days at 29°C, flipping onto freshly prepared food every 2 days. The sterility of GF flies was confirmed by plating fly homogenate on MRS plates 1 day prior to *Vc* infection.

### Generation of germ-free fish

Adult TL zebrafish were bred for less than 60 minutes to minimize exposure to microbes from parents. Embryos were collected, washed, and split into GF or CR cohorts. The GF cohort was kept in sterile EM supplemented with antibiotics (100µg/mL Ampicillin, 5µg/mL Kanamycin, 250ng/mL Amphotericin B, and 5µg/mL Gentamicin), while the CR cohort was kept in EM. Embryos were incubated at 29°C for 4-6 hours and washed every 2 hours with EM or EM plus antibiotics for CR and GF cohorts respectively. Once embryos were at 50% epiboly, the GF cohort was washed three times with sterile EM, and then 2 minutes with 0.1% Polyvinylpyrrolidone-iodine (PVP-I) in EM. Embryos were rinsed three times with EM and then immersed in 0.003% sodium hypochlorite (bleach) solution for 20 minutes. GF embryos were washed three more times and transferred into tissue culture flasks with sterile EM. CR embryos received the same number and duration of washes with sterile EM rather than PVP-I or bleach. GF confirmation was performed at 4 days post fertilization by plating out 100 µL EM from flasks onto TSA plates. Parental tank water and sterile EM were used as positive and negative control respectively, where bacterial growth was confirmed in tank water and absent in sterile EM. GF and CR flasks with bacterial absent and present respectively were used for subsequent analysis.

### *Vc* oral infection

Virgin female flies were kept at 29°C for 7 days prior to infection. *esg^ts^>shn^RNAi^* flies and their wild-type counterparts (*esg^ts^/+*) were maintained at 18°C for 10 days and then shifted to 29°C for 3 days to minimize detrimental effects of prolonged BMP inactivation. For oral infection in flies, *Vc* C6706 and C6706Δ*vasK*^47^ were grown on LB plates (0.5% NaCl, 0.5% yeast extract, 1% tryptone, 1.5% agar) supplemented with 100 μg/ml streptomycin (Sigma SLBK5521V) for 16-18 hours at 37°C. To prepare a *Vc* infection culture, single colonies were removed from the plate, suspended in LB broth, and diluted to a final OD600 of 0.125. Flies were starved for 2 hours at 29°C prior to infection. In each vial, 10-15 flies were placed onto one-third of a cotton plug soaked with 3 ml of LB broth (Uninfected) or *Vc* infection culture (C6706 or C6706Δ*vasK*). Vials were kept at 29°C with a 12 hour/12 hour light/ dark cycle for oral infection.

For *Vc* infection in zebrafish, a single colony of C6706 or C6706Δ*vasK* was suspended in LB broth with 100 μg/ml streptomycin and grown with aeration overnight at 37°C. *Vc* cells were washed twice with PBS and diluted to OD600 of 1 (∼10^8^ CFU/mL). To infect fish larvae, 15-20 larvae (6 days post fertilization) were incubated in each well of a 6-well plate containing 4 mL embryo medium with 20 μL of OD600 = 1 infection culture (C6706 or C6706Δ*vasK*) or 20 μL PBS (Uninfected) for 24 hours. For infection in adult fish, 5 female fish per treatment were fasted for ∼20 hours and then incubated in a 400 mL beaker containing 200 mL filter sterilized fish tank water with 1 mL OD600 = 1 culture (C6706 or C6706Δ*vasK*) or 1 mL PBS (Uninfected) for 24 hours. Both larvae and adult fish were kept in a 29°C incubator with a 14-hour/10-hour light/ dark cycle during infection.

### Immunofluorescence

Fly intestines were dissected in PBS, fixed in 8% formaldehyde in PBS for 20 minutes, washed in PBS with 0.2% Triton-X (PBST) for 30 minutes, and blocked in PBST with 3% bovine serum albumin (BSA) for 1 hour at room temperature. Guts were stained overnight at 4°C in PBST + 3% BSA with appropriate primary antibodies. The following day guts were washed in PBST and stained for 1 hour at room temperature with secondary antibodies and DNA stain (1/1000 Hoechst 33258; Invitrogen H3569) in PBST + 3% BSA. Guts were washed in PBST for 30 minutes at room temperature then in PBS overnight at 4°C.

Zebrafish larvae (6 day-post-fertilization) were incubated in embryo medium with 1% DMSO and 5 mM EdU (Invitrogen C10340) for 8 hours at 29°C after the 16 hour *Vc* infection. Fish intestines were dissected in PBS and fixed in 4% paraformaldehyde in PBS overnight at 4°C. Guts were washed three times with PBSTx (0.75% TritonX-100 and 0.02% NaN_3_ in PBS), blocked for 1 hour in PBSTx + 3% BSA at room temperature, and stained with primary antibodies in blocking buffer overnight at 4°C. Guts were washed with PBSTx and then stained for 1 hour at room temperature with secondary antibodies and nuclear stain (1/1000 Hoechst 33258; Invitrogen H3569), followed by rinse in PBSTx. EdU detection was performed by incubating guts in Click-iT^®^ reaction cocktail (Invitrogen C10340) for 30 minutes at room temperature. Guts were washed in PBSTx followed by extra washing in PBS. Whole fly guts and fish larval intestines were mounted on microscope slides using Flouromount^TM^ (Sigma; F4680) and visualized with a spinning-disk confocal microscope (Olympus IX-81 motorised microscope base with Yokagawa CSU 10 spinning-disk scan-head).

Primary antibody used in this study are chicken anti-GFP (1/2000; Invitrogen PA1-9533), rabbit anti-PH3 (1/1000; Milipore 06-570), rabbit anti-pSmad3 (1/200; Abcam ab52903), mouse anti-armadillo (1/100; DSHB N2 7A1), mouse anti-Delta (1/100; DSHB C594.9B), and mouse anti-perospero (1/100; DSHB MR1A). Secondary antibodies used are goat anti-chicken 488 (Invitrogen A11039), goat anti-rabbit 568 (Invitrogen A11011) and goat anti-mouse 647 (Invitrogen A21235) at a concentration of 1/1000.

### Immunohistochemistry

Intestines from adult zebrafish were dissected and fixed in BT fixative (0.15 mM CaCl_2_, 0.1 M PO_4_, 4% sucrose, 4% paraformaldehyde in dH_2_O) for 48 hours at 4°C. Guts were embedded in paraffin, sectioned into 5 μm slices, and collected on Superfrost Plus slides. Sections were deparaffinized with Toluene and rehydrated. Antigen unmasking was performed by boiling slides in 0.1 M sodium citrate buffer (pH 6) in a 98°C water bath for 20 minutes. Sections were incubated in 3% hydrogen peroxide, washed with PBSt (0.5% TritonX-100 in PBS), blocked with 10% normal goat serum in PBSt, and incubated with mouse anti-PCNA (1/20000; Sigma P8825) overnight at 4°C in a humid chamber. Sections were washed three times with PBSt and incubated in SignalStain^®^ Boost Detection Reagent (HRP, Mouse; CST 8125P) for 30 minutes at room temperature. SignalStain^®^ DAB Chromogen solution (CST 8059) was added onto each section for colorimetric detection. Slides were rinsed and counterstained with ¼-strength hematoxylin. All sections were dehydrated and mounted in Dpx. Images were captured using ZEISS AXIO A1 compound light microscope with SeBaCam 5.1MP camera, and qualifications were done using Fiji software.

### Transmission electron microscopy

Fly posterior midguts were cut into 1 mm pieces and immediately placed into fixative (3% paraformaldehyde and 3% glutaraldehyde in 0.1 M cacodylate buffer with 0.1 M CaCl_2_, pH 7.2). Tissue processing was performed at the Cell Imaging Facility at University of Alberta. The midgut sagittal sections from 5 flies per treatment were visualized using a JEOL 2100 transmission electron microscope.

### Quantification of cells per gut area

Fly posterior midgut (R4/R5) was located by identifying the midgut-hindgut transition area and moving 1-2 frames anterior from the attachment site of the Malpighian tubules. Z-slice images of fly posterior midguts and entire zebrafish larval guts were acquired using Perkin Elmer’s Volocity software. Cells in each image frame were manually quantified in Fiji software. PH3+ cells in the entire fly midgut were counted through the eyepiece of the microscope. All guts damaged by the dissection process were excluded from quantification. Data points in the same figure were results from one single experiment. Finally, the quantification of cells from Figures 2 and 7 was reanalyzed in a double-blinded study to confirm the findings.

### Statistical analysis and figure construction

All graphs and plots were constructed using R (version 4.1.1) via R-studio (version 2021.09.0-315) with easyGgplot2 (version 1.0.0.9000). One-way Analysis of Variance (ANOVA) was used to determine the overall statistical difference, a Tukey’s test for Honest Significant Differences was used for multiple comparisons, and an unpaired student t-test was used to compare two groups. Details of the specific test used for each data panel can be found in the tables and figure captions. Statistical significance was set at p ≤ 0.05. Figures in were assembled using Inkscape.

## SUPPLEMENTARY FIGURES

**Supplementary Figure 1.**
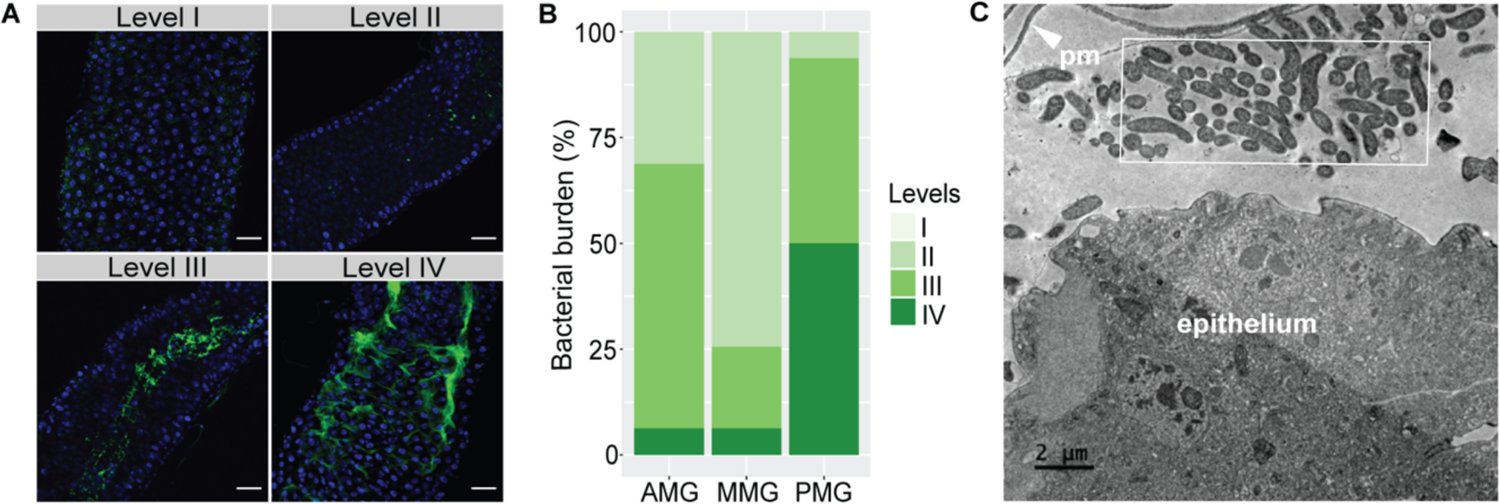
*Vc* colonizes the posterior midgut of *Drosophila*. **(A)** Immunofluorescence of fly midguts infected with C6706. DNA stained by Hoechst (blue) and GFP-labeled C6706 (green). Four different levels of bacterial burden: level I = no visible GFP-labeled bacteria, level II = a few scattered individual GFP-C6706, and large clumps of *Vc* in the lumen (level III) or in close proximity to the intestinal epithelium (level IV). Scale bars = 25 μm. **(B)** Approximate bacterial burden in different regions of fly guts infected with C6706 (n=20) for 24 h. Assessment based on the four-level scale in (A). Anterior midgut, AMG; middle midgut, MMG; posterior midgut, PMG. **(C)** Transmission electron microscopy of the posterior midguts of flies infected with C6706 for 24 h. Clusters of rod-shaped *Vc* are indicated with a white box. Peritrophic matrix, pm.

**Supplementary Figure 2.**
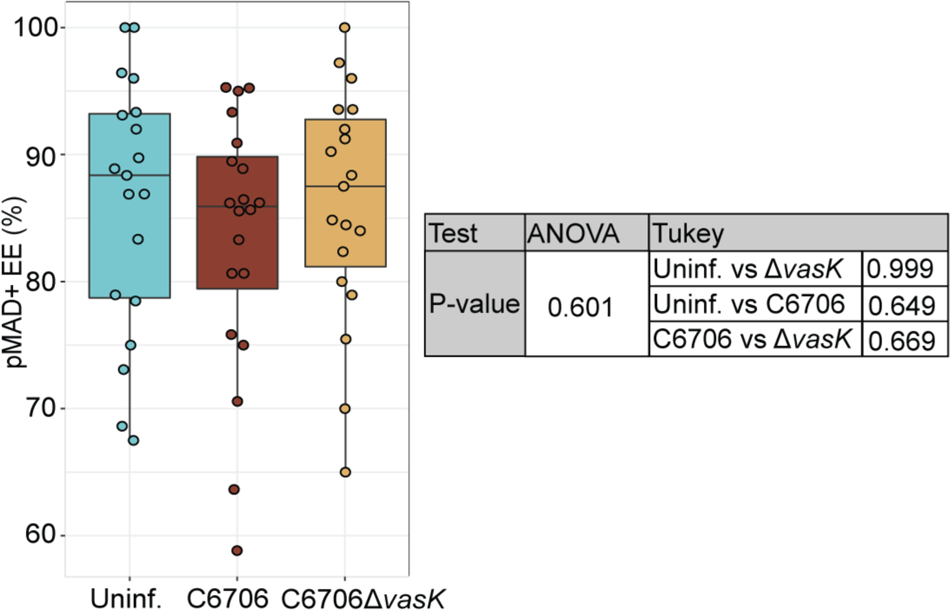
BMP activation in enteroendocrine cells. Percentage of Pros+ enteroendocrine cells (EE) that are pMAD+ in uninfected *esg^ts^*/+ guts, and guts infected with C6706Δ*vasK* or C6706. Each dot represents a measurement from a single fly gut. P values were calculated using the significance tests indicated in the table.

**Supplementary Figure 3.**
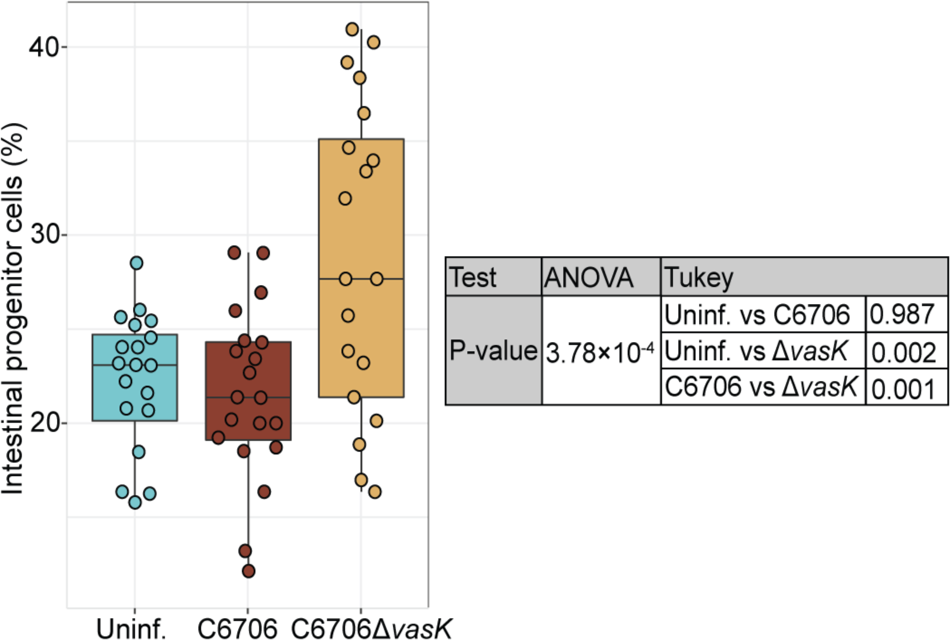
*Vc* T6SS limits progenitor expansion. Percentages of progenitor cells in uninfected, C6706Δ*vasK*-infected or C6706-infected *esg^ts^*/+ guts. Each dot represents a measurement from a single fly. P values were calculated using the significance tests indicated in the table.

**Supplementary Figure 4.**
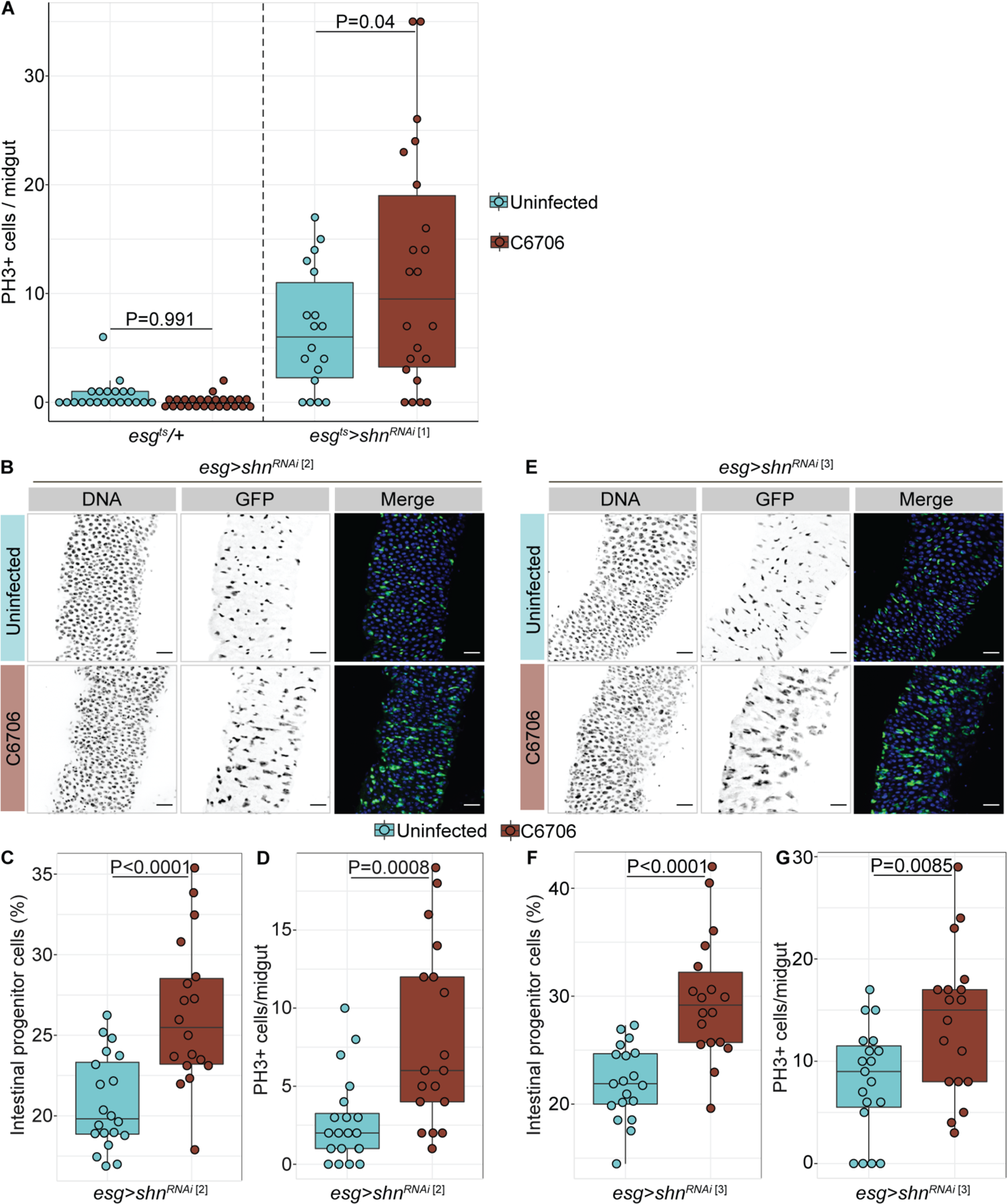
BMP regulates cell proliferation after *Vc* infection. Cell proliferation and epithelial repair were measured in flies with progenitor-specific shn knockdown by independent RNAi lines upon infection with C6706 for 36h. **(A)** PH3+ cell per midgut in wild-type (*esg^ts^/+*) or *esg^ts^>shn^RNAi^*^[1]^ flies. **(B-D) (B)** images of posterior midguts, **(C)** percentages of GFP+ progenitors, and **(D)** numbers of PH3+ cells in uninfected and C6706-infected *esg^ts^>shn^RNAi^* ^[2]^ flies. **(E-G) (E)** images of posterior midguts, **(F)** percentages of GFP+ progenitors, and **(G)** numbers of PH3+ cells in uninfected and C6706-infected *esg^ts^>shn^RNAi^* ^[3]^ flies. Each dot represents a measurement from a single fly gut. P values are calculated using unpaired Student t-tests.

